# Using a bistable animal opsin for switchable and scalable optogenetic inhibition of neurons

**DOI:** 10.1101/2020.06.03.129627

**Authors:** Jessica Rodgers, Beatriz Bano-Otalora, Mino DC Belle, Sarika Paul, Rebecca Hughes, Phillip Wright, Richard McDowell, Nina Milosavljevic, Patrycja Orlowska-Feuer, Franck Martial, Jonathan Wynne, Edward R Ballister, Riccardo Storchi, Annette E Allen, Timothy Brown, Robert J Lucas

**Affiliations:** Faculty of Biology Medicine and Health, University of Manchester, Manchester, UK; Institute of Biomedical and Clinical Sciences, University of Exeter Medical School, University of Exeter, Exeter, UK; Department of Neurophysiology and Chronobiology, Institute of Zoology and Biomedical Research, Jagiellonian University in Krakow, Krakow, Poland; Department of Biomedical Engineering, Columbia University, New York, USA

**Keywords:** Optogenetics, Opsins, GPCR, Neuronal Inhibition, Neuronal Silencing, Bistable, light-activated G-protein-coupled receptors

## Abstract

There is no consensus on the best optogenetic tool for neuronal inhibition. Lamprey parapinopsin (‘Lamplight’) is a Gi/o-coupled bistable animal opsin that can be activated and deactivated by short and long wavelength light, respectively. Since native mechanisms of neuronal inhibition frequently employ Gi/o signalling, we asked here whether Lamplight could be used for optogenetic silencing. We show that short (405nm) and long (525nm) wavelength pulses repeatedly switch Lamplight between stable signalling active and inactive states, and that combining these wavelengths can be used to achieve intermediate levels of activity. We demonstrate that these properties can be applied to produce switchable and scalable neuronal hyperpolarisation, and suppression of spontaneous spike firing in the mouse hypothalamic suprachiasmatic nucleus. We show that expressing Lamplight in (predominantly) ON bipolar cells can photosensitise retinas following advanced photoreceptor degeneration, and that 405 and 525nm stimuli can produce responses of opposite sign in output neurons of the retina. Lamplight-driven responses to both activating (405nm) and deactivating (525nm) light can occur within 500ms and be elicited by intensities at least 10x below threshold for available inhibitory optogenetic tools. We conclude that Lamplight can co-opt endogenous signalling mechanisms to allow optogenetic inhibition that is scalable, sustained and rapidly reversible.

## Introduction

Inhibitory optogenetic tools are light sensitive proteins or opsins which can be expressed in neurons to cause hyperpolarisation and silencing of spike firing ^1,2^. Although several inhibitory optogenetic tools have been developed, there is no consensus regarding the best to use for either basic research or therapeutic applications. Early inhibitory optogenetic tools, such as the inwardly directed chloride pump Halorhodopsin ^3,4^ and the outwardly directed proton pump Archaerhodopsin ^5^, provide hyperpolarisation with millisecond temporal resolution, but also require very bright light. Later generation tools, including the red-shifted chloride pump JAWS ^6^ and anion conducting channelrhodopsins, such as gtACR ^7^ are sensitive to lower levels of light across a wider range of wavelengths, but require repeated light stimulation to maintain neuronal inhibition. Step function tools, the chloride-conducting channelrhodopsin SwiChR ^8^ and the light-gated potassium channel Blink2 ^9,10^, have recently been developed for sustained inhibition over the second-minute timescale. However, tools based upon light-gated ion channels or pumps have the potential to cause unpredictable and abnormal cellular responses, including non-physiological ion concentration gradients and antidromic spiking ^11,12^, and, in some cases, unexpected excitatory responses to light ^13,14^. A more physiological alternative would be to take advantage of endogenous cellular processes capable of causing neuronal inhibition, such as Gi/o signalling. Gi/o proteins can activate G-protein coupled inwardly rectifying potassium (GIRK) channels to induce hyperpolarisation ^15,16^. This is a widespread mechanism, lying downstream of native GPCRs providing inhibitory neuronal modulation, and co-opted by the inhibitory chemogenetic tool, h4MDi ^17,18^. Gi/o signalling is also involved in GIRK-independent synaptic silencing, inhibiting presynaptic neurotransmitter release ^19,20^

Optogenetic control over Gi/o-protein signalling could be provided by the family of animal opsins, which function as light sensitive G-protein coupled receptors (GPCRs). Indeed, naturally occurring Gi/o coupled opsins, such as Rod opsin, LWS and SWS cone opsin and the engineered Gi-coupled OptoXR (a mu-Opioid receptor-Rod opsin chimera), have already been used as optogenetic tools, mostly as a method of restoring visual responses in the degenerate retina ^21–23^, but also to produce neural inhibition in the brain ^24,25^. However, these rod and cone opsin-derived tools have a fundamental characteristic that limits the degree to which their activity can be controlled under heterologous expression: while they are activated by light, their deactivation relies on light-independent mechanisms whose activity is largely outside the experimenter’s control. This means that although they may be switched on by light, more subtle control over the degree of activation and its longevity is hard to achieve. The natural diversity of animal opsins provides an attractive potential solution to this problem. Several classes of animal opsins are thermally stable in both photo-activated and ‘dark’ states and are thus termed ‘bistable’. Critically, bistable opsins may be both switched on and off by light, raising the possibility of achieving close control over the timing and degree of G-protein signalling.

Here, we set out to determine whether bistable animal opsins could form the basis of switchable inhibitory optogenetic tools. For this purpose, we used the lamprey parapineal opsin, parapinopsin (termed here Lamplight). Lamplight represents an attractive candidate for optogenetic control because it couples to Gi *in vitro* ^26^, and forms active and inactive states with very different wavelength selectivity (peak sensitivity (λ_max_ = 370nm and 515nm, respectively ^26–29^). Here we show that these characteristics allow switchable and scalable inhibition of neurons using appropriate application of short and long wavelength light at safer light intensities.

## Results

### Lamplight allows switchable, titratable control of Go activity

We first set out to explore Lamplight’s potential for achieving switchable control over G-protein signalling under heterologous expression. For this purpose we adapted a live cell bioluminescent resonance energy transfer (BRET) assay to report activation of Go (the most abundantly expressed member of the Giot family ^30^). In brief, Hek293T cells were transiently co-transfected with plasmids driving expression of Lamplight, Gαo, Gβ and Gγ subunits tagged with a components of a split fluorescent protein, and a G-protein receptor kinase (GRK3) fragment tagged with nanoluciferase (**Figure 1A**). In this way, Lamplight-driven activation of Go may be revealed as a light dependent increase in BRET ratio associated with interaction of the GRK3 fragment with liberated Gβ/γ dimer ^31^. This approach resulted in Lamplight expression in a subset of Hek293T cells, correctly localised to the plasma membrane (**Figure 1A**), and a clear increase in BRET signal in response to a 1s flash of 405nm light (**Figure 1B**). These data confirm that photoactivated Lamplight can liberate Go.

**Figure 1.**
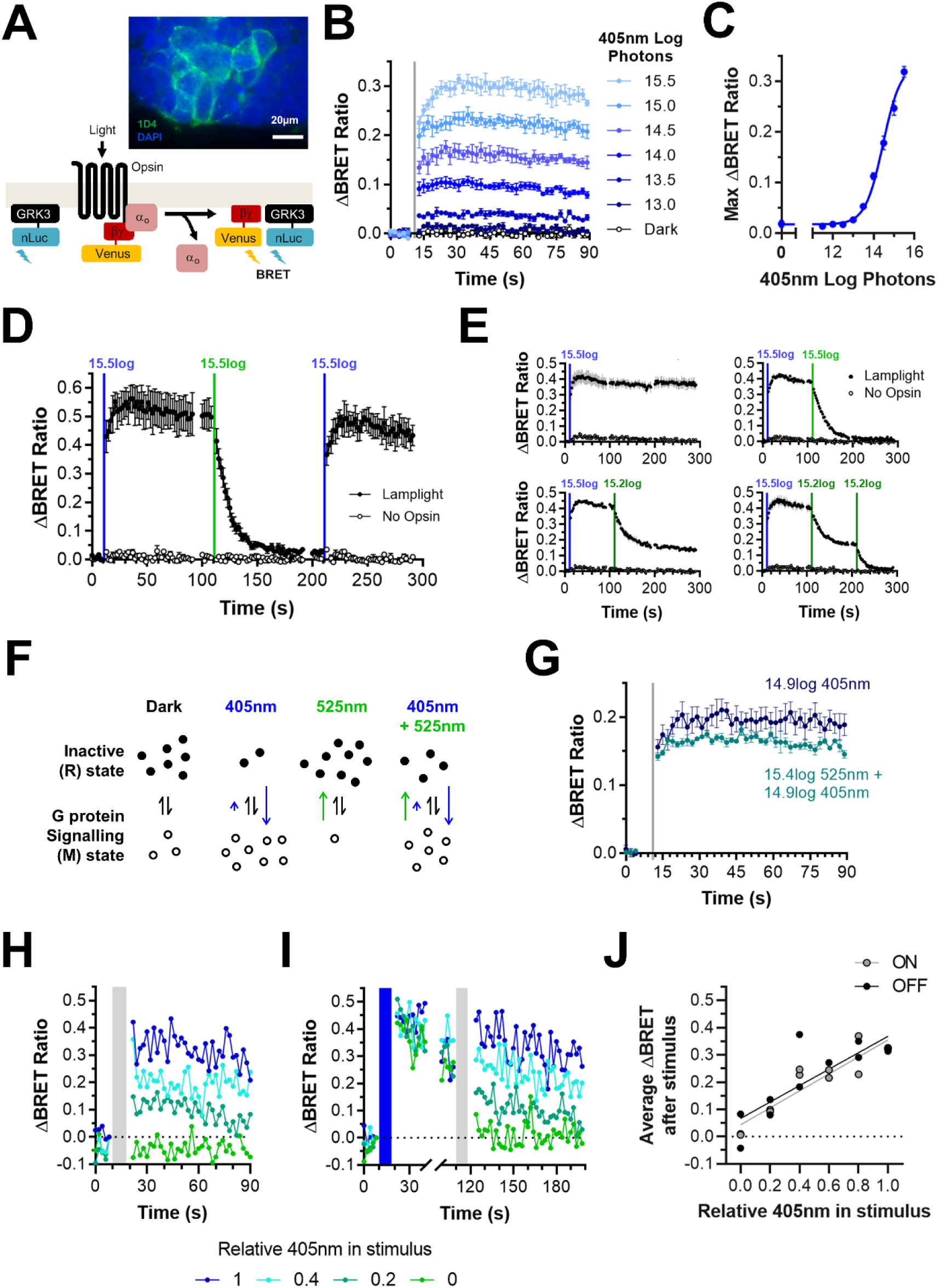
Lamplight can be used for switchable and tuneable control Go activity in Hek293T cells. **A)** Immunocytochemistry of Hek293T cells transiently transfected with Lamplight-1D4 plasmid (anti-1D4 in green, DAPI in blue) with, below, schematic of BRET assay, which measures interaction between opsin-liberated Gβγ dimer (tagged with fluorescent Venus) and GRK3 fragment tagged with nanoluciferase (bottom). **B)** Time course of Lamplight-driven response to 1s flashes of increasing intensity 405nm light (grey line). **C)** Lamplight 405nm irradiance response curve. **D)** Time course of G protein activity in response to 405nm (blue line) and 525nm light (green line) for Lamplight (filled circles) and negative control (unfilled circles). **E**) Time course of Lamplight response to sequential presentation of 405nm (blue line) followed by 525nm light of different intensities (green). **F)** Schematic of Lamplight photoequilibrium. In dark, Lamplight is mostly in inactive state (filled circles). Exposure to 405nm light shifts equilibrium (blue arrows) so opsin is mostly in the signalling state (unfilled circles). Exposure to 525nm light shifts equilibrium (green arrow) so Lamplight is pushed into the inactive state. Presenting 525nm and 405nm light together results in small increase of opsin in signalling state due to opposite direction shifts partially cancelling each other. Ratio of symbols and arrow size are not quantitative. **G)** Lamplight response to 405nm light (dark blue) can be antagonised by simultaneous presentation of 525nm light (light blue). **H-I)** Representative time courses of Lamplight-driven responses to 8s mixed stimuli (grey bar) with different 405nm: 525nm ratios from 1 (100% 405nm) to 0 (100% 525nm) from **H)** dark or **I)** after 8s 405nm light (blue bar). **J)** Sustained Lamplight-driven G protein activity (measured as average BRET ratio from 20-80s after stimulus) after exposure to 8s mixed stimuli from dark (ON, filled circles) or 405nm light-adapted state (OFF, unfilled circles). Fit with linear trendline (Best fitting parameters for ON, slope = 0.311, Y-intercept = 0.04, R^2^ = 0.85. For OFF, slope = 0.30, Y-intercept = 0.07, R^2^ = 0.69). For all graphs, error bars show SEM, N = 3-4 replicates. All intensities are for total Lamplight effective log photons/cm^2^/s.

Animal opsins frequently drive responses at lower light intensities than required by other optogenetic actuators, such as microbial opsins. Altering the intensity of the 405nm pulse modulated the BRET signal amplitude across a wide range of intensities, from 13-16 log photons/cm^2^/s (**Figure 1B**). Lamplight sensitivity (log EC50 = 14.4log photons/cm^2^/s, **Figure 1C**) was broadly consistent with other animal opsins we have tested using BRET assay (Rod opsin log EC50 for 480nm = 14.6log photons or 1.5µW/mm^2^), around 1000-fold less light than required by Halorhodopsin and Archaerhodopsin under heterologous expression (17-18log photons/cm^2^/s or 1-10mW/mm^2^) ^3–5^. The threshold for a measurable response was 13.5 log photons/cm^2^/s, but note that signal amplification at later steps in the G-protein signalling cascade could render integrated cellular responses substantially more sensitive.

We next set out to determine how closely Go activity in this system could be controlled using Lamplight. The BRET signal in Lamplight expressing cells remained elevated over tens of seconds following 1s 405nm illumination, suggesting continuous opsin signalling (as expected for generation of a thermally stable active photoproduct). We therefore asked if presentation of a light pulse predicted to be optimally absorbed by Lamplight’s activated state was able to switch Go activity off. Indeed, a 1s 525nm flash resulted in relaxation of the BRET signal back to baseline (**Figure 1D**). Note that, although this relaxation occurred over several seconds, this time course likely reflects the time taken for decay of the GRK3:beta-gamma complex, as light-induced changes in opsin conformation are near instantaneous. These characteristics suggest that Lamplight could function as a switchable optogenetic tool in this environment, and indeed a subsequent 405nm pulse switched the system back ‘ON’, producing a sustained high BRET signal (**Figure 1D**).

Lamplight thus allows switchable control over Go signalling. We next sought to establish the degree to which it could provide quantitative control over the level of Go activation. Our initial experiments showed that different intensities of 405nm could be used to scale the amplitude of the Lamplight response (**Figure 1B**). In theory a similar effect should be possible starting from an activated state by modulating the amount of subsequent 525nm exposure. This proved to be the case. Thus, while a single 405nm pulse established a high BRET signal that could be fully switched off with a subsequent bright 525nm pulse (**Figure 1E**, top), a lower intensity 525nm pulse established an intermediate BRET signal that was sustained for the remainder of the recording (**Figure 1E**, bottom left). Moreover, sequential exposures to the lower intensity 525nm produced stepwise reductions in BRET signal (**Figure 1E**, bottom right).

An alternative approach to tuning signal activity for bistable opsins can be to adjust the ratio of activating vs deactivating wavelengths in incident light. Thus, under extended exposure, bistable opsins are expected to reach an equilibrium at which fractional concentrations of active and inactive states are defined by the spectral composition of the light. In the case of Lamplight, stimuli rich in short wavelengths should produce a pool strongly biased toward the active state, while stimuli rich in long wavelengths should enrich the inactive state (**Figure 1F**). In such a system, controlling the ratio of short to long wavelength light can provide quantitative control over G-protein signalling that is not dependent upon the starting state concentrations of the two opsin states (and thus on the pattern of prior light exposure). We therefore set out to determine whether Lamplight showed such behaviour in this system. Our first step was to ascertain whether inclusion of 525nm antagonised the effects of a 405nm pulse in dark adapted cells. This proved to be the case (**Figure 1G**). We next presented mixtures of 405 and 525nm light at a range of ratios (see methods) to dark adapted cells and to cells previously photoactivated by 405nm (**Figure 1H&I**). The duration of light exposure was increased to 8s for these 405+525nm mixtures to maximise the possibility that the system had approached equilibrium. We found that indeed the BRET signal magnitude was impacted by the wavelength ratio, with larger responses elicited by stimuli biased in favour of 405nm (**Figure 1H&I**). This occurred in both dark-adapted and previously photoactivated cells, and indeed, these bispectral stimuli produced equivalent BRET signals in cells under these two different initial conditions (**Figure 1J**). These data indicate that tuning the ratio of short vs long wavelength light achieves quantitative control over Lamplight signalling independent of previous light exposure.

Our findings in Hek293T cells suggest that Lamplight can be used for switchable (step function) control of Go activity. Lamplight also allows quantitative control over G-protein signalling using two approaches: 1.) changing the intensity of the activating or deactivating stimuli; or 2.) varying the spectral composition of incident light stimulus.

### Lamplight as an inhibitory optogenetic tool for neuronal silencing

Having established Lamplight can be used for switchable control of Go activity, we turned to the question of whether this could be applied to neuronal control. We expressed Lamplight in neurons of the mouse hypothalamus using viral gene delivery. Double floxed inverse orientation (DIO) Lamplight-T2A-mCherry AAV was injected bilaterally into the suprachiasmatic nucleus (SCN) of *VIP*^*Cre/+*^ mice (which express Cre-recombinase in a subset of SCN neurons). VIP-expressing cells, like other neurons in the mouse SCN, spontaneously fire at higher rates during the circadian light phase ^32–37^, making them suitable for exploring inhibitory effects of Lamplight. Moreover, as these neurons are not readily silenced by a commonly applied channel-based optogenetic inhibitor (ArchT)^36^, they represent an opportunity to trial Lamplight in a challenging cell type. At 4 weeks post-injection, we made targeted whole cell current clamp recordings from mCherry fluorescent neurons in living SCN slices (**Figure 2A**).

**Figure 2.**
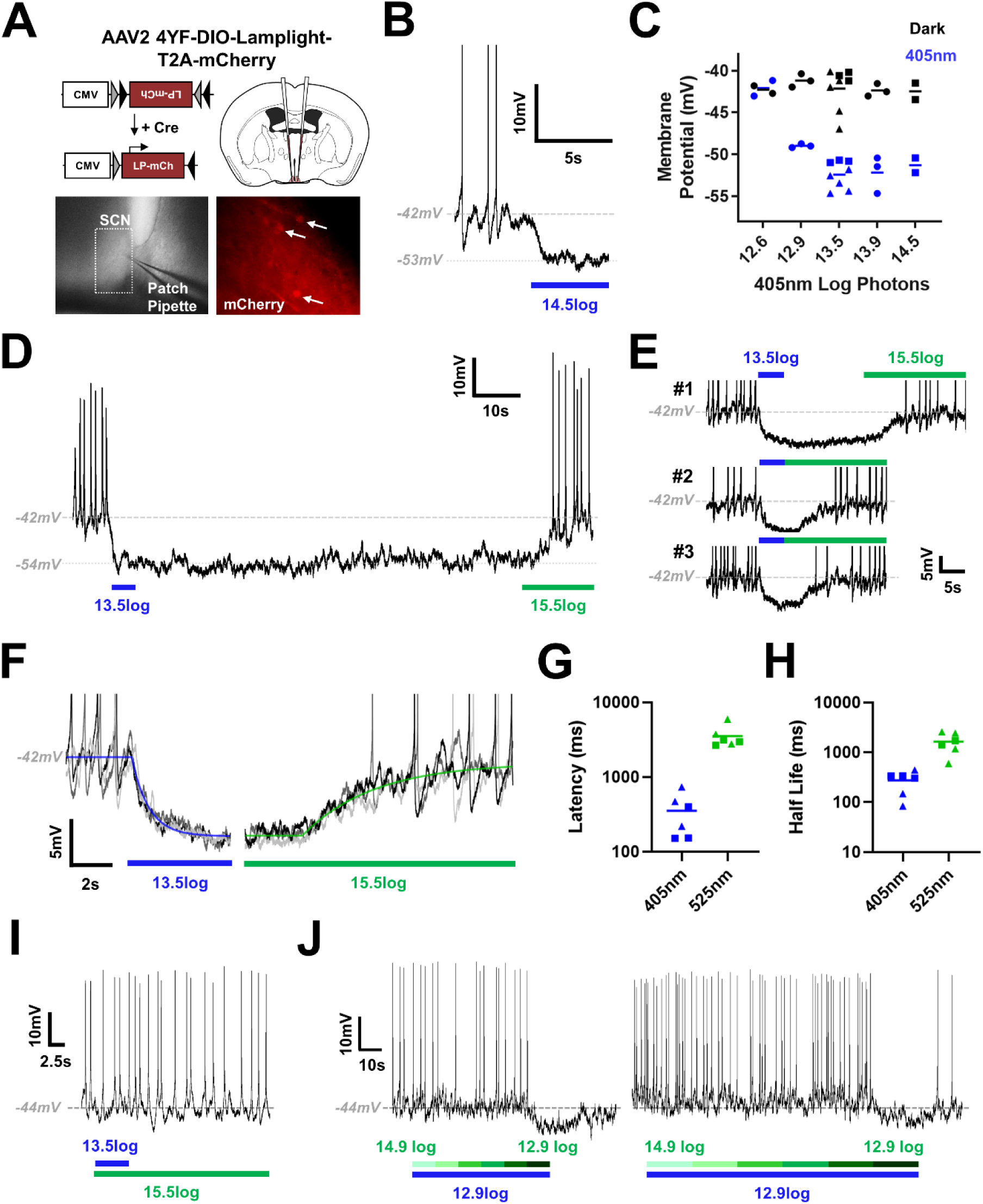
Lamplight causes sustained reversible hyperpolarisation in SCN neurons. **A)** Schematic showing bilateral delivery of Lamplight AAV bilaterally to the suprachiasmatic nucleus (SCN) of *VIP*^*Cre/+*^ mice. Whole cell patching recordings on coronal SCN slices (bottom left) were made by targeting mCherry fluorescent neurons (bottom right) within the SCN. **B)** Representative trace showing hyperpolarisation in a Lamplight-expressing SCN neuron after exposure to 405nm light (blue bar). **C)** Hyperpolarisation of resting membrane potential to 405nm (blue symbols) compared to dark baseline (black symbols) was observed over a range of intensities. Recordings were made from 3 different cells across 2 SCN slices. **D)** Representative trace showing hyperpolarisation and silencing of neuronal firing were sustained during dark after exposure to 405nm light (5s, blue bar). This inhibition can be reversed by subsequent exposure to 525nm light (15s, green bar). **E)** Hyperpolarisation and depolarisation of Lamplight-expressing neurons by 405nm (blue bar) and 525nm (green bar), respectively, could be reliably observed across multiple stimulus presentations. **F)** Three example traces from a single cell exposed to 405nm (13.5log photons) followed by 525nm (15.5log photons/cm^2^/s). Individual traces were fit with plateau followed by exponential decay (405nm, blue) or association curves (525nm, green) using non-linear regression for 6 trials across 2 cells. Parameters from best-fit curves were used to calculate **G)** latency (ms) and **H)** half-life (ms). **I)** Simultaneous presentation of 525nm (green bar) and 405nm (blue bar) prevents hyperpolarisation of neuron and inhibition of spike firing. **J)** Constant 405nm light (12.9log photons/cm^2^/s, blue bar) was presented with 525nm light (green bar) of decreasing intensity from 14.9log to 12.9log photons/cm^2^/s in 6 steps of either 10s (left) or 20s (right). 525nm light antagonised the hyperpolarising effects of 405nm light for most intensities except 12.9log photons/cm^2^/s. For **C), G)** and **H)** data from different cells is indicated by different shaped symbols. For **B), E)** and **F)** action potentials are truncated for clarity.

Short wavelength light (405nm) caused hyperpolarisation and suppression of spike firing in mCherry-labelled cells (**Figure 2B**). We recorded this inhibitory response for a range of different 405nm light intensities from 14.5 to 12.9 log photons/cm^2^/s (**Figure 2C**), with mean change in resting membrane potential (RMP) from −42.1±0.5mV in dark to −51.6±0.5 mV at the end of the 5s 405nm stimulus (mean±SEM). We were unable to detect changes in RMP at 12.6log photons/cm^2^/s, suggesting that 12.9log photons/cm^2^/s is close to threshold for opsin-driven activity.

Lamplight-induced silencing had two particularly interesting properties. First, we found that hyperpolarisation was sustained, with new RMP remaining stable once the 405nm light was switched off (**Figure 2D**). Second, in the majority of mCherry fluorescent cells tested, we found that this inhibition could be reversed by subsequent presentation of 525nm light (**Figure 2D-G**), with average RMP of −44.1±0.5mV (mean±SEM) after 15s of 525nm light (although some cells did not respond to the longer wavelength **Supplementary Figure 1**). For cells where the inhibition caused by 405nm could be switched off with 525nm light, we observed this switching behaviour reliably across repeated stimulus presentations (**Figure 2E**).

We next focused on the kinetics of responses to 405nm and subsequent 525nm light. Using non-linear regression, individual traces of membrane potential were fit with exponential decay or association curves to aid estimation of response latency and velocity (**Figure 2F**). Onset of hyperpolarisation occurred rapidly to 405nm with mean latency of 353.3±92.6ms and half-life of 273.9±54.2ms (**Figure 2G-H**, mean±SEM). In comparison, it took several seconds for depolarisation by 525nm to begin, with mean latency of 3.5±0.5s, and responses were much slower, with average half-life of 1.6±0.3s (mean±SEM).

Finally, we examined how neurons responded to simultaneous presentation of blue and green light. 525nm light was able to antagonise the hyperpolarisation caused by 405nm light when presented concurrently. No change in RMP was observed when these two wavelengths were used simultaneously (**Figure 2I**), −43.5±0.6mV, compared to preceding dark baseline, - 43.2±0.9mV (mean±SEM). We found 15.5 log photons/cm^2^/s of 525nm light was sufficient to inhibit responses to both 13.5 and 14.5 log photons/cm^2^/s 405nm light, suggesting Lamplight expressed in neurons can also be controlled by modulating the spectral composition of light. To explore this further, we presented dim blue light (12.9 log photons/cm^2^/s) and superimposed green light of decreasing intensity across 6 steps (14.9-12.9 log photons/cm^2^/s). For both 10s and 20s steps, hyperpolarisation could be observed for dimmest 525nm light (12.9log photons/cm^2^/s) – suggesting varying ratio of 405nm and 525nm light will lead to different degrees of hyperpolarisation (**Figure 2J**) and that Lamplight can be used as a tuneable inhibitory optogenetic tool to control neuronal firing.

### Using Lamplight to restore visual responses to the degenerate retina

The SCN recordings confirm that Lamplight may be used to switchable control of neuronal activity. In theory, such switching could have high temporal resolution, as light-induced changes in opsin state are near instantaneous. However, the actual spatiotemporal resolution of optogenetic control provided by Lamplight is predicted to be additionally influenced by the rate at which downstream elements of the G-protein cascade are activated and deactivated. Indeed, the return to baseline activity following 525nm occurred over several seconds in SCN neurons. We then finally applied Lamplight to a system in which temporal fidelity is a more critical feature of control. To this end, we turned to the vertebrate retina, a system in which G-protein signalling is known to have high temporal resolution to gain a better appreciation of its potential performance. In the intact retina, visual signals are conveyed by G-protein signalling cascades at two stages – within the photoreceptor itself, and at the synapse between photoreceptors and ON bipolar cells. The ON bipolar cell response relies upon a Go signalling cascade ^38^ and so represents an attractive potential target for Lamplight control.

We introduced the DIO-AAV2-4YF-Lamplight-T2A-mCherry virus to the vitreal space of a transgenic mouse (*L7*^*Cre*^) driving expression of floxed transgenes in rod (ON) bipolar cells and a small subset of retinal ganglion cells (RGCs) ^39^. To eliminate conventional visual responses, we employed animals homozygous for the *Pde6b*^*rd1*^ mutation, which causes rapid and extensive degeneration of rod and cone photoreceptors ^40,41^. As expected, we found transgene expression in the inner nuclear layer (location of ON bipolar cells) and a few RGCs in *L7*^*Cre*^; *Pde6b*^*rd1/rd1*^ mice treated in this way (**Figure 3A**).

**Figure 3.**
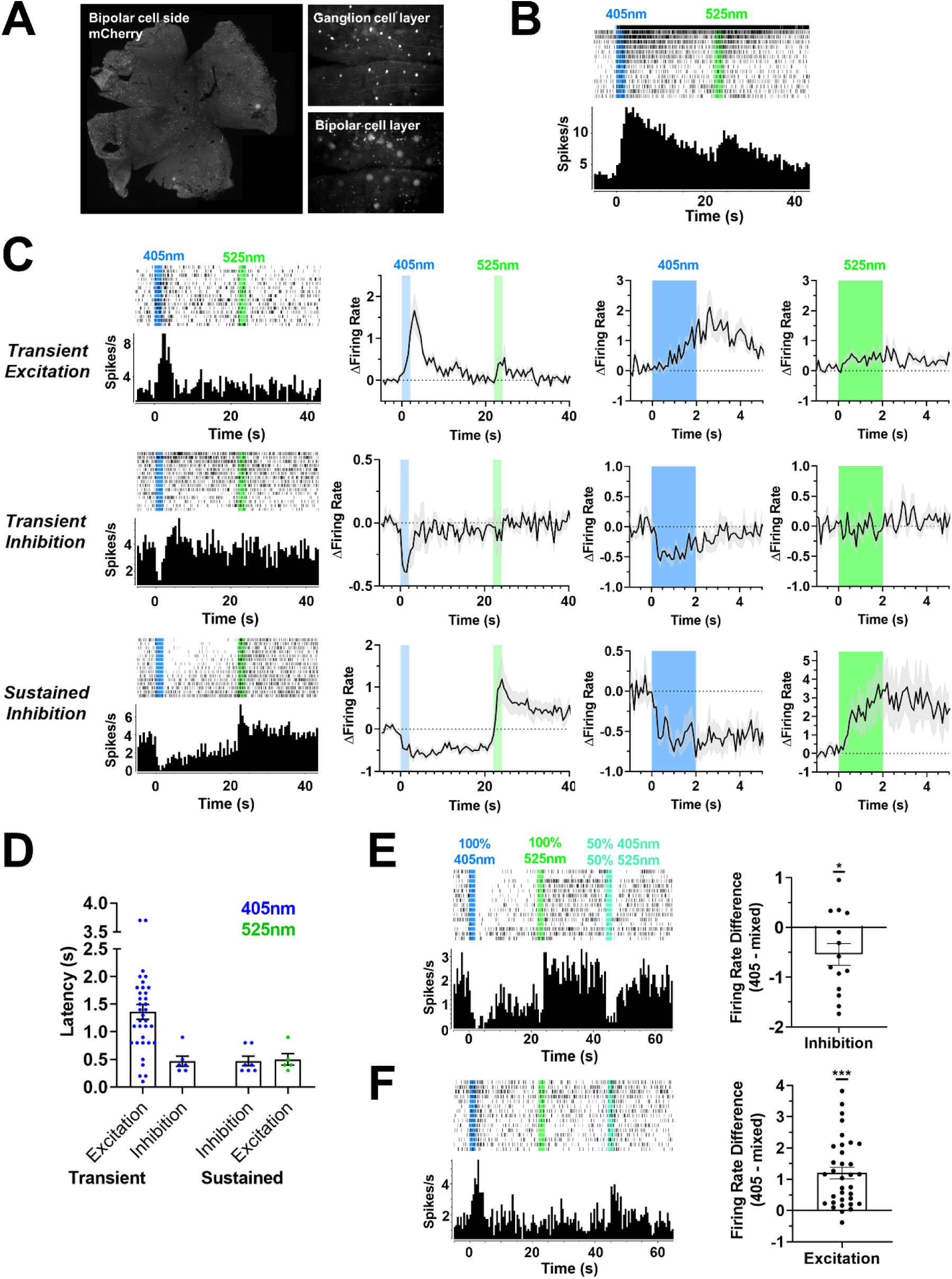
Lamplight expression in degenerate retinas can restore diverse and switchable responses to light. **A)** Immunohistochemistry of *L7Cre rd1* retina after intravitreal injection of Lamplight-T2A-mCherry virus. Anti-mCherry staining (white) was widespread (Left). Fluorescent cell bodies were found in different focal planes consistent with expression in ganglion (Top Right) and bipolar cell layer (Bottom Right). **B)** Representative single unit showing characteristic ipRGC response. **C)** Three categories of Lamplight-driven responses were identified based on response to 2s 405nm light across 15 trials – transient excitation (top, N = 35), transient inhibition (middle, N = 6), and sustained inhibition (bottom, N = 8). Left panel shows representative single units from each category. Centre left panel shows mean baseline normalised firing rate (grey shaded area shows SEM, 500ms bins). Centre and far right panels show baseline normalised firing rate during 405nm and 525nm light stimuli, respectively (100ms bins, N numbers as above, except N = 5 for sustained response to 525nm). **D)** Distribution of response latencies for 3 categories of Lamplight-driven responses to 405nm, as well as reversal caused by 525nm in sustained inhibition units. Two outliers with slow latency (>2.5s) in transient excitation group are likely responding to stimulus offset. **E)** Inhibition responses were smaller and less sustained for mixed stimuli (50% 405nm, 50% 525nm) compared to 405nm only. **F)** Excitation responses were attenuated for mixed stimuli compared to 405nm only. Left panel shows representative retinal single unit. For **E-F)** Left panel shows representative single unit. Right panel shows distribution of difference in firing rate for 405nm vs mixed stimulus. For inhibition, this was calculated as difference in average firing rate across interstimulus interval. For excitation, this was calculated as difference in average firing rate in first 5s from stimulus onset. All representative single units show perievent rasters (first trial at top) and associated perievent firing rate histograms (Bin size = 500ms). Timing of light stimuli shown by shaded vertical bars. All light stimuli are 16 log total Lamplight effective photons/cm^2^/s. Error bars show SEM. * p<0.05, *** p < 0.001 (One-sample t-test compared to zero).

To assess the light response of Lamplight-expressing retinas, we placed excised retinas, RGC side-down onto a multi-electrode array. In this way we were able to record the spiking activity of RGCs (which represents the output signal of the retina) in response to light pulses impacting Lamplight expressed in the inner retina. Based upon the known spectral sensitivity of Lamplight, and our own experience in other cell types, we presented transduced retinas with sequential 2s flashes of 405nm and 525nm (20s interstimulus interval). Leaving aside persistent increases in firing to both wavelengths (**Figure 3B**), which could be produced by surviving melanopsin photoreceptors ^42–44^, we observed three qualitatively distinct classes of response to this stimulus (**Figure 3C)**. The most abundant was an increase in firing to 405nm, typically lasting a few seconds (**Figure 3C**, top panel, N = 35 units from 4 retinas). A subset of RGCs with this excitatory response also showed a significant increase in firing also to 525nm, but others were insensitive to the long wavelength stimulus. The other two classes of 405nm-response were both inhibitory but differed in temporal characteristics (**Figure 3C** middle and bottom panels). In one (**Figure 3C**, middle panel), 405nm induced a transient reduction in firing (N = 6 units from 4 retinas), lasting little longer than the period of light exposure. RGCs with such transient inhibition did not show any clear response to 525nm. In the other (**Figure 3C**, bottom panel), the inhibition of firing induced by 405nm was persistent, lasting many seconds after the light pulse had terminated (N = 7 units from 3 retinas). This final response class most faithfully recreated the switching behaviour of Lamplight observed in other systems, as the persistent reduction in firing induced by 405nm was accompanied by an equally reproducible increase in firing in response to 525nm. In the RGCs displaying this property, firing rate could thus be switched between low and high states by sequential presentations of 405 and 525nm light. The diversity of RGC responses to light in the Lamplight retina is consistent with the established ability of the retinal circuitry to invert polarity and apply different temporal frequency filters during signal transfer, and with the outcome of other studies restoring photosensitivity to ON bipolar cells ^21–23^.

A more detailed examination of response profiles (**Figure 3C**, right) revealed that all 3 response types started soon after the onset of light exposure. The slowest in this regard were the transient excitations to 405nm, which on average built up over several seconds and had a mean response latency of 1.22s (range 0.1 – 3.7s, **Figure 3D**). By contrast, inhibitory responses were more immediate (Figure 3C) with response latencies of 0.47s (range 0.3-0.9s) and 0.47s (range 0.3–0.8s) for transient and sustained responses respectively; **Figure 3D**). Excitatory responses to 525nm were also relatively rapid (Figure 3D), with mean response latency of 0.50s (range 0.3 – 0.9s, **Figure 3D**).

We finally asked if it were possible to control the amplitude of Lamplight responses in the retina using the approaches employed in SCN neurons and HEK293 cells. To this end, we compared responses to 405nm light alone with those elicited by a 50:50 mix of 405nm and 525nm light. We found that, indeed the mixed light produced a smaller response in cells with all 3 response classes (**Figure 3E-F**). Thus, the degree of reduction in firing was smaller in cells with either transient or sustained inhibition to 405nm (mean±SEM difference in response = 0.54±0.22 spikes/s; One-sample t-test compared to 0, t(13) = 2.47, p = 0.028, **Figure 3E**), as was the increase in firing in cells excited by 405nm (mean ± SEM difference in response = 1.2±0.19spike/s; One-sample t-test compared to 0, t(33) = 6.47, p < 0.001, **Figure 3F**).

## Discussion

Here, we have explored the potential of Lamplight, a bistable light-sensitive GPCR, as an inhibitory optogenetic tool. We have demonstrated that Lamplight can be activated and inactivated using exposure to short and long wavelength light (respectively) at low intensities to produce reversible hyperpolarisation in the brain and photo-switchable changes in the retina. In both *in vivo* systems, single light pulses could produce sustained inhibition of spike firing, with onset occurring within 500ms. The bistable nature of Lamplight allows its degree of activity to be controlled by modulating either the intensity or spectral composition of the light stimulus. Our data establish bistable animal opsins as promising high sensitivity optogenetic tools for switchable neuronal inhibition.

Lamplight compares favourably with currently available inhibitory optogenetic tools in several regards. First, it is sensitive to relatively low levels of light. In SCN neurons we were able to detect responses down to 1s of 12.9log photons/cm^2^/s 405nm light (equivalent to 0.15uW/mm^2^). That makes Lamplight 10,000-fold more sensitive than first generation inhibitory optogenetic tools, such as halorhodopsin ^3,4^ and archaerhodopsin ^5^, which require light stimuli of ∼10mW/mm^2^. Lamplight is still ∼100-fold more sensitive for 1s flashes than microbial opsins engineered for improved light sensitivity, such as gtACR2 ^7^, to 1s flashes of light. This differential becomes even more marked for sustained inhibition, which required repeated stimulation of gtACR2 but not Lamplight. Even available step function tools developed for sustained inhibition, such as Blink2 ^9,10^ are about 10-fold less sensitive than Lamplight, and often require brighter or longer duration stimuli. The high sensitivity of Lamplight reduces both the potential for neuronal inhibition to be accompanied by damaging levels of light and heat, and the challenge of controlling a widely dispersed population of neurons.

Lamplight’s ability to provide sustained and switchable inhibition compares favourably with available alternatives. In all systems we tested, Lamplight-driven effects on activity could be measured for tens of seconds after a 405nm pulse and could be subsequently switched off by 525nm light. Such prolonged inhibition is not practicable with most inhibitory actuators, which would require constant or repeated illumination with high intensity light. Some switchable tools have been developed, such as the bistable anion channel SwiChR ^8^, which can be used for inhibition over similar timescales to Lamplight, albeit with reduced light sensitivity. Another functionality afforded by Lamplight’s bistability is the ability to control the degree of G-protein activity by modulating the spectral composition of light. Our experiments confirm that altering the ratio of short to long wavelength light can be used to titrate the light response. Such ‘ratiometric’ control could be a useful complement to the standard method of controlling opsin activation by altering light intensity, as it has the potential to provide a more uniform level of activity across a population of neurons differing in the amount of opsin expressed, and lying at different distances from a light source.

Lamplight targets endogenous mechanisms of neuronal inhibition and could be widely used for experimental, and potentially therapeutic, applications. Gi/o signalling is a ubiquitous cellular phenomenon ^45^, with Go widely expressed in the central nervous system (CNS)^30^. Numerous native receptors inhibit neuronal activity using Gi/o mechanisms, including opioid receptors involved in pain perception and analgesia, as well as dopamine receptors important for learning and reward ^46^. CNS disorders are amongst the most abundant targets for GPCR drug discovery ^47^, with GPCRs that couple to Gi/o involved in conditions including schizophrenia, Parkinson’s disease, and Alzheimer’s disease. While it is possible that Lamplight may not be able to hyperpolarise neurons without Gi/o-coupled GIRK channels, work using Gi/o-coupled chemogenetic tools has demonstrated that so far this has not been a concern ^48^. In future, chimera between lamplight and non-photosensitive GPCRs may be applied to tune its G-protein specificity and perhaps more appropriately engage native receptor signalling ^49^.

The important role of Go signalling in visual transduction at the synapse between photoreceptor and ON bipolar cell ^38,50^ also makes Lamplight a potential candidate for vision restoration therapy. We have shown here that expressing Lamplight in, predominantly, ON bipolar cells restores photosensitivity in a murine model of advanced retinal degeneration. Lamplight activation in the inner retina supported rapid increases and decreases in firing of the retina’s output neurons (RGCs). The switchable nature of Lamplight control was also apparent in a subset of RGCs in which 405 and 525nm stimuli drove antagonistic changes in ganglion cell firing. These characteristics suggest that Lamplight’s ability to generate sustained and rapidly tuneable Go activation could be helpful in recreating the sort of high fidelity, analogue input signal that the retina is able to process to support high spatiotemporal resolution vision.

Of the many bistable animal opsins described, we chose Lamplight for these studies because it has several characteristics that make it well suited for optogenetic inhibition. Firstly, it is Gi/o coupled, allowing access to endogenous mechanisms of neuronal hyperpolarisation and synaptic inhibition. Secondly, it has a clear separation in spectral sensitivity between its thermally stable inactive and active states. This feature is critical in allowing different wavelengths of light to activate and inactivate the pigment. Thirdly, the spectral sensitivity of its active state is long wavelength shifted compared to that of the inactive state. Opsin photopigments have a stereotyped spectral sensitivity profile that is characterised by a much more complete loss of sensitivity at wavelengths longer vs shorter than the peak. For this reason, differences in relative sensitivity between the 2 pigment states can be much greater at longer wavelengths of light in comparison to shorter wavelengths. In the case of an optogenetic tool, if one chooses a pigment in which the spectral sensitivity of the active state is long-wavelength shifted compared to that of the inactive state, long wavelength light can be used very effectively to deplete the amount of pigment in the active state. Conversely, although short wavelengths will increase the fractional concentration of inactive state in pigments whose active state is short-wavelength shifted, a substantial fraction of pigment will remain in the active state. Achieving more complete inactivation has the benefit of being able both to fully silence the optogenetic tool and allow much greater fractional increases in activity. Perhaps the biggest disadvantage to the choice of Lamplight is that it is most effectively activated by UV light (λ_max_ = 370nm). We show here that this does not preclude activation by >400nm light at intensities at least 10-fold lower than other switchable inhibitory optogenetic tools, but a pigment shifted to longer wavelengths could enhance sensitivity and allow control by a greater range of visible wavelengths.

In conclusion, Lamplight is an efficient inhibitory optogenetic tool that can be used for sustained and reversible neuronal silencing. It allows optogenetic control that is physiological and can be tuned by altering intensity and spectral composition of light stimulus. The unique properties of Lamplight make it well suited to basic research of GPCR signalling biology as well as therapeutic applications.

## Methods

### Cell Culture

Hek293T cells (ATCC) were incubated at 37°C (5% CO_2_) in culture media (Dulbecco’s modified Eagle’s medium with 4500mg/L glucose, L-glutamine, sodium pyruvate and sodium bicarbonate from Sigma) with penicillin (100U/ml), streptomycin (100μ/ml) and 10% fetal bovine serum (FBS). Cell were seeded into 12-well plates at a density of 250 000 cells/well in antibiotic-free culture medium. After 48hrs, cells were transiently transfected using Lipofectamine 2000 (Thermo Fisher) according to manufacturer’s instructions.

For BRET G protein activation assay, each well of 12-well plate was transfected with following: 100ng sVβ1, 100ng sVγ2, 25ng mGRK3-nLuc, 50ng Gαo and 500ng opsin (where applicable). BRET assay plasmids were obtained from Kiril Martemyanov & Ikuo Masuho (Scripps Research Institute). All subsequent steps were conducted under dim red light. After addition of transfection reagent and DNA, cells were incubated for 4-6hours at 37°C, then resuspended in 1ml of culture media containing 10μM 9-*cis* retinal. 100μl of cell suspension was added to each well of a white-walled clear-bottomed 96-well plate (Greiner Bio-One) and left overnight before performing BRET G protein activation assay.

For immunocytochemistry, cells were plated onto coverslips and transfected with plasmid as described above. After 24hrs, cells were fixed using 4% paraformaldehyde (PFA) in PBS and washed with PBS. All dilutions were carried out in PBS with 0.05% Tween-20. Cells were first blocked in 5% normal donkey serum, then incubated in 1:500 dilution of primary antibody (mouse anti-1D4, Abcam, catalogue number ab5417) with 1% donkey serum for 1hr at room temperature. Cells were washed 3 times, then incubated in 1:500 dilution of secondary antibody (Donkey anti-mouse 488, Abcam, catalogue number A-210202) for 30mins at room temperature in the dark. Cells were washed 4 times, then mounted onto slides using Prolong Anti-fade Gold media with DAPI and allowed to dry overnight at room temperature in dark.

### BRET G protein activation assay

All following steps were carried out under dim red light. Before beginning the BRET G protein activation assay, culture media was removed from cells and replaced with 50μl imaging media (L-15 media without Phenol Red containing L-glutamine (Gibco), 1% FBS and 10μM 9-*cis* retinal). Cells were then left to incubate at room temperature in dark for up to 2hours.Under dim red light, NanoGlo Live Cell substrate (Furimazine derivative, Promega) was diluted 1:40 in PBS. 12.5ul of dilute NanoGlo substrate solution was added to each well of 96-well plate (final dilution of 1:200), for up 6 wells at a time, and incubated for 5mins to allow luminescence to equilibrate. BRET measurements were conducted using a FluoStar Optima microplate reader (BMG Labtech). As this plate reader has a single photomultiplier tube (PMT), light emitted by fluorescent Venus and bioluminescent Nanoluc were measured sequentially using 535nm (30nm FWHM with gain set to 4095) and 470nm (30nm FWHM with gain set to 3600) emission filters. A 0.68s recording interval was used for each filter, with a total cycle time of 2s. We coupled the plate plater bottom optic to the liquid light guide of a pE-4000 light source (CoolLED) to allow us to flash cells inside the plate. To avoid bleaching the PMT during light stimulus, a motorized shutter was built to protect the PMT while light was on. The activity of this shutter was synced to the light source using an Arduino microcontroller. To avoid neighbouring wells being exposed to light, each recorded well was surrounded by empty wells and the location of wells measured was counterbalanced. For every recording, we measured 5 cycles of baseline measurement (10s total) and then flashed cells with 1s of light before resuming recording for up to 45cycles (90s total). For repeated light exposure, sequential recordings of the same well were performed. Timing of light stimuli and recording were controlled using plate reader software (Optima in script mode). A range of different light stimuli were used (all 1s in duration), described in more detail in **Table S1**. The spectral power distribution of the stimuli was measured using a SpectroCAL MKII spectroradiometer (Cambridge Research Systems Ltd).

### Animals

All experiments were conducted in accordance with the UK Animals (Scientific Procedures) Act (1986). *L7*^*Cre/+* 51^ *rd1* transgenic mice on a mixed C3H x C57Bl/6 background were used for retinal MEA experiments. L7Cre mice express Cre recombinase in rod bipolar cells and a subset of retinal ganglion cells ^39^. They also possess the *Pde6b*^*rd1*^ mutation ^40,41^ causes progressive retinal degeneration, with vision loss complete once animals are over 80 days old. *L7Cre rd1* mice were genotyped to confirm they do not possess the GRP179 point mutation ^52^ which affects bipolar cell function. *VIP*^*Cre/+*^ mice ^53^ on a mixed C57BL/6 × 129S4Sv/Jae background were used for SCN patch electrophysiology experiments. These mice possess Cre recombinase in vasoactive intestinal polypeptide (VIP)-expressing cells. All mice were kept under a 12:12 light dark cycle with food and water provided *ad libitum*.

### AAV virus

*L7*^*Cre/+*^ *rd1* mice received bilateral intravitreal injections of lamprey parapinopsin, termed Lamplight, virus (AAV2 4YF– ITR - DIO-CMV-LPPN-1D4-T2A-mCherry – WPRE-SV40 late polyA - ITR). The Lamplight virus was packaged in an AAV2/2 capsid with four tyrosine to phenylalanine mutations ^54^ to achieve efficient viral transduction of retinal cells, in particular bipolar cells. An 8 amino-acid C-1D4 tag (ETSQVAPA) was added to the C-terminus of the lamprey parapinopsin transgene (AB116380.1). The Lamprey parapinopsin-1D4 transgene (LPPN-1D4) was linked to a mCherry fluorescent reporter using a T2A sequence to ensure 1:1 co-expression of the two proteins. The inverted LPPN-1D4-T2A-mCherry open reading frame was flanked by two pairs of Lox sites (LoxP and Lox2272), so that in the presence of Cre recombinase, the transgene is inverted into the sense orientation and expression is driven by the constitutive CMV (cytomegalovirus) promotor. A woodchuck hepatitis virus post-transcriptional regulatory element (WPRE) and SV40 late polyA sequence were also included between ITRs to improve transgene expression. Virus was obtained from VectorBuilder.

### Brain injections

Heterozygous *VIP*^*Cre/+*^ mice were injected aged 10-11 weeks old using previously established protocol^36^. Briefly, mice were anaesthetised using 1% isoflurane and positioned in a stereotaxic frame. Craniotomy and bilateral AAV injection was carried out using a computer-controlled motorised injector (Drill and 473 Microinjection Robot, Neurostar) and 69nl of Lamplight-T2A-mCherry virus was injected into SCN at 25nl/s. Micropipettes were left in place for 5mins after injection, then removed in 100µm/5s increments until outside SCN.

Mice were culled by cervical dislocation during the light phase (ZT4-8) 6 weeks after SCN injection. Brain slices were prepared as described previously^55,56^. Briefly, brains were removed and mounted on metal stage. Coronal slices containing mid-SCN levels (250uM) across the rostro-caudal axis were cut using a Campden 7000smz-2 vibrating microtome (Camden Instruments). Slicing was in ice-cold (4°C) sucrose-based incubation solution containing: 3mM KCl, 1.25mM NaH_2_PO_4_, 0.1mM CaCl_2_, 5mM MgSO_4_, 26mM NaHCO_3_, 10mM D-glucose, 189mM sucrose, oxygenated with 95% O_2_, 5%CO_2_. After cutting, slices were left to recover at room temperature in a holding chamber with continuously gassed incubation solution for at least 20 min before transferring into artificial cerebrospinal fluid (aCSF). aCSF composition was the following: 124mM NaCl, 3mM KCl, 24mM NaHCO_3_, 1.25mM NaH_2_PO_4_, 1mM MgSO_4_, 10mM D-Glucose, 2mM CaCl_2_, and 0mM sucrose.

### Whole-cell current clamp recordings

Coronal brain slices containing mid-level of the SCN were placed in the bath chamber of an upright Leica epi-fluorescence microscope (DMLFS; Leica Microsystems Ltd) equipped with infra-red video-enhanced differential interference contrast (IR/DIC) optics. Slices were held in place using a tissue anchor grid and continuously perfused with aCSF by gravity (∼2.5ml/min) without addition of 9-*cis* retinal. Brightfield photographs of the patch pipette sealed to these neurons were taken at the end of each recording for accurate confirmation of the anatomical location of the recorded cell within the SCN.

Patch pipettes (resistance 7–10MΩ) were fashioned from thick-walled borosilicate glass capillaries (Harvard Apparatus) pulled using a two-stage micropipette puller (PB-10). Recording pipettes were filled with an intracellular solution containing the following: 130mM K-gluconate, 10mM KCl, 2mM MgCl_2_, 2mM K2-ATP, 0.5mM Na-GTP, 10mM HEPES, and 0.5mM EGTA, pH adjusted to 7.3 with KOH, measured osmolarity 295–300 mOsmol/kg). Cell membrane was ruptured under minimal holding negative pressure.

An Axopatch Multiclamp 700A amplifier (Molecular Devices) was used in a current-clamp mode with no holding current (I=0), after establishing whole cell configuration at −70 mV. Signals were sampled at 25 kHz and acquired in gap-free mode using pClamp 10.7 (Molecular Devices). Access resistance for the cells used for analysis was <30 MΩ. All data acquisition and protocols were generated through a Digidata 1322A interface (Molecular Devices). All recordings were performed at room temperature (∼ 23°C).

mCherry fluorescent SCN neurons were identified with a 40x water immersion UV objective (HCX APO; Leica) using a cooled Teledyne Photometrics camera (Retiga Electro). Red fluorescence was detected using 550nm excitation light from pE-4000 light source (CoolLED) and a Leica N2.1 filter cube (Dichroic = 580nm, Emission filter = LP 590nm). The same light source was used to deliver 405nm and 525nm light stimuli via the 40x UV objective. Patching data were analysed using Clamplit 10.7, Prism and Excel. Resting membrane potential was measured manually in pClamp for 10s before stimulus and at end of 405nm stimulus. For kinetics analysis, data were first downsampled (extract every 100^th^ data point, new sample frequency = 4ms). For responses to 405nm, data from first 3s after stimulus onset were fit with a plateau followed by either an exponential decay or exponential association curve. For responses to 525nm, we used data from first 15s after stimulus onset. We excluded any data >1 standard deviation from mean (to remove spikes that might distort fit), then fit with plateau followed by exponential association. Non-linear curve fitting was performed using least-squares minimisation in Excel. Data from 2 replicates were excluded due to poor curve fits (R^2^ < 0.3).

### Intravitreal Injections

Heterozygous *L7*^*Cre/+*^ *rd1* mice were injected aged 9-10 weeks. Mice were anaesthetised by intraperitoneal injection ketamine (75mg/kg body weight) and medetomidine (1mg/kg body weight). Once anaesthetised, mice were positioned on a heat mat to prevent cooling. Pupils were dilated with 1% tropicamide eye drops (Bausch & Lomb) and a 13mm coverslip was positioned on gel lubricant (Lubrithal) applied to the cornea. Approximately 2.2μl of virus (1.12 × 10^13^ genomic counts per ml) was injected into the vitreous of each eye using a Nanofil 10μl syringe (World Precision Instruments) using 35-gauge bevelled needle using a surgical microscope (M620 F20, Leica). All mice received bilateral injections. Anaesthesia was reversed by intraperitoneal injection of atipamezole (3mg/kg body weight). During recovery, 0.5% bupivacaine hydrochloride and 0.5% chloramphenicol was applied topically to the injected eyes. Mice also received 0.25ml of warm saline by subcutaneous injection to aid recovery.

### Multi-electrode array recordings

Multi-electrode array recordings were conducted between 13 to 15 weeks after bilateral intraocular injection. Mice were dark adapted overnight. All following steps were performed under diffuse dim red light. Dark adapted mice were culled by cervical dislocation (approved Schedule 1 method). Enucleated eyes were placed in petri dish filled with carboxygenated (95% O_2_/ 5% CO_2_) Ames’ media (supplemented with 1.9g/L sodium bicarbonate, pH 7.4, Sigma Aldrich) and retinas dissected, with care taken to remove vitreous from inner retinal surface. Retinal wholemounts were then placed on glass coated metal harps (ALA Scientific Instruments) and positioned ganglion-cell side down on coated 256-channel multi-electrode arrays (MEA, Multi Channel Systems).

Multi-electrode arrays were first incubated in fetal bovine serum overnight at 4°C, then coated with 0.1% polyethyleneimine (PEI) in borate buffer (pH8.4) for 1hr at room temperature. PEI coating was then removed, and MEA washed 4-6 times with ddH_2_0. PEI-coated MEAs were then air-dried and coated with 20μg/ml laminin in fresh Ames’ medium for 30-45mins at room temperature ^57,58^. Laminin solution was removed before retina was positioned on the MEA. Once in place on the MEA, the retina was continuously perfused with carboxygenated Ames’ media with 10μM 9-*cis* retinal at 2-3ml/min using a peristaltic pump (PPS2, Multi Channel Systems) and maintained at 34°C using a water bath heater (36 °C), in-line perfusion heater (35 °C) and base plate heater (34 °C). Once positioned, retinas were perfused in dark for at least 45mins before first light stimuli were applied.

Data were sampled at 25kHz using MC Rack software (Multi Channel Systems). A Butterworth 200Hz high pass filter was applied to raw electrode data to remove low frequency noise. Amplitude threshold for spike detection was 4-4.5standard deviations from baseline. Light stimuli were presented using a customised light engine (Thorlab LEDs). An Arduino Due microcontroller controlled by programmes written in LabVIEW (National Instruments) to control stimulus duration and intensity by altering LED output. **Table S1** shows the intensity of light stimuli used for testing Lamplight activity.

Multi-unit data were spike sorted into single units using Plexon Offline Sorter (using Template Sorting Method and Principal Component Analysis, followed by manual verification), then analysed using Neuroexplorer, Matlab 2012 and Prism. Single units with light responses were identified using following approach. A perievent stimulus time histogram (PSTH) was generated for first 15 trials with 500ms bin size. Lamplight-driven light responses were identified as significant change in spike firing rate within 5s of 405nm stimulus onset – defined as 2st deviations from mean baseline (5s before 405nm stimulus onset) for at least 2 bins (1s). Units were excluded if they had insufficient spiking to provide meaningful baseline spiking rate (defined as spiking in <50% baseline bins), had square wave shape caused by removal of noisy spikes during spike sorting, or responses that were not reproducible across trials (15 trials were split into groups of 3, then PSTH compared to confirm reproducibility of responses). The remaining 52 units were then divided into categories based on their response profile. For latency, we used data in 100ms bins and identified first bin after light stimulus where data was more than 2standard deviation from baseline (5s before stimulus onset). For comparing responses from mixed and 405nm only stimuli, we used data in 500ms bins. For transient excitation responses, we compared average spike firing rate for first 5s after stimulus onset. For inhibition responses, we compared average spike firing rate across interstimulus interval (20s total).

### Immunohistochemistry

Injected mice were culled by cervical dislocation (approved schedule 1 method). Eyes were removed and placed in 4% paraformaldehyde in phosphate buffered saline for 24hrs at 4°C. Retinas were then dissected and permeabilised in 1% Triton-X in PBS for 3x 10mins at room temperature. Retinas were then blocked using 1% Triton-X in PBS with 10% normal goat serum for 2-3 hours while shaking gently. Retinas were incubated in primary antibody (1:500 dilution of rabbit anti-mCherry antibody, Kerafast, catalogue no. EMU106, in 1% Triton-X in PBS with 2.5% goat serum) overnight at room temperature. Retinas were washed in PBS with 0.2% Triton-X for 4 × 30mins shaking gently. Retinas were incubated in secondary antibody (Goat anti-rabbit Alexa fluorophore 555, Abcam, catalogue no. ab150078, in 1% Triton-X in PBS with 2.5% goat serum) for 3-4 hours at room temperature in the dark. Retinas were washed a further 4 times in PBS with 0.2% Triton-X for 30mins each, then washed in ddH_2_0 for 10mins. Retina was then mounted on a microscope slide using Prolong Gold anti-fade mountant and allowed to dry overnight at room temperature.

### Fluorescence microscopy

Images were acquired using a Leica DM2500 microscope with DFC365 FX camera (Leica) and a CoolLED-pE300-white light source. Imaging software was Leica Application Suite Advanced Fluorescence6000. Images were acquired using Chroma ET Y3 (excitation = 545nm, emission = 610nm), A4 (excitation = 360nm, emission = 470nm), and L5 (excitation = 480nm, emission = 527nm) filter sets. Global enhancements to image brightness and contrast were made using ImageJ software.

### Generating ratiometric bispectral light stimuli

Spectrum of Monochromatic light stimuli (405nm and 525nm) were measured using a SpectroCAL MKII spectroradiometer (Cambridge Research Systems, Ltd) from 380-750nm Spectra were converted from W/m^2^ to Photons/cm^2^/s. Log effective photons for inactive state and active state for each wavelength were calculated by multiplying light spectra by log relative sensitivity of opsin at each wavelength. Opsin nomograms for Lamplight inactive and active state were generated with λmax of 370nm and 515nm ^28^, respectively, using A1-visual pigment template ^59^. In addition to the effective photons for the inactive or active state, the total effective photons for Lamplight (ie: light that either state could respond to) were calculated by summing effective photons for both states. Throughout text, unless specified otherwise, intensities provided are in total Lamplight effective photons.

To generate ratiometric stimuli, the total Lamplight effective photons for 405nm only or 525nm only stimuli were first matched. Take 15.5log total Lamplight effective photons/cm^2^/s. For the 405nm only stimulus, this would result in 15.29log inactive state effective photons and 15.09log active state effective photons, 60% of total effective photons are for inactive state, while remaining 40% are for active state. For the 525nm only stimulus, 15.5log total Lamplight effective photons, would mean 12.18log inactive effective photons/cm^2^/s and 15.5 log active state effective photons/cm^2^/s, 100% of total effective photons are for active state, with 0% for inactive state. These matched values are then used to define ratiometric bispectral stimuli on a scale of 1 to 0 relative amount of 405nm light in the stimulus (where 1 = 405nm light only or maximum possible R effective photons, and 0 = 525nm light only minimum possible inactive state effective photons). Continuing the example above, for 15.5log total Lamplight effective photons/cm^2^/s, a stimulus with 0.25 relative 405nm light in stimulus, would require 14.68log inactive state effective photons/cm^2^/s and 15.43log active state effective photons/cm^2^/s, where 0.15% of total effective photons are for inactive state and 85% of total effective photons are for active state (0.15/0.6 = 0.25 of maximum inactive state effective photons).

## Acknowledgements

This work was funded by grants from Human Frontier Science Program (RGP0034/2014) and Medical Research Council (MR/N012992/1 and MR/S026266/1) awarded to RJL. POF was funded by the Bekker Programme implemented by the Polish National Agency for Academic Exchange.

## Author contributions

Experiments were designed by JR, RJL, NM, EB and AA. Data were acquired and analysed by JR, BBO, MB, SP, RH, PW, RM, NM, POF, FM, JW, RS, AA, and TB. Manuscript was written by JR and RJL. All authors approved the submitted version.

## Figure Legends

**Figure S1.**
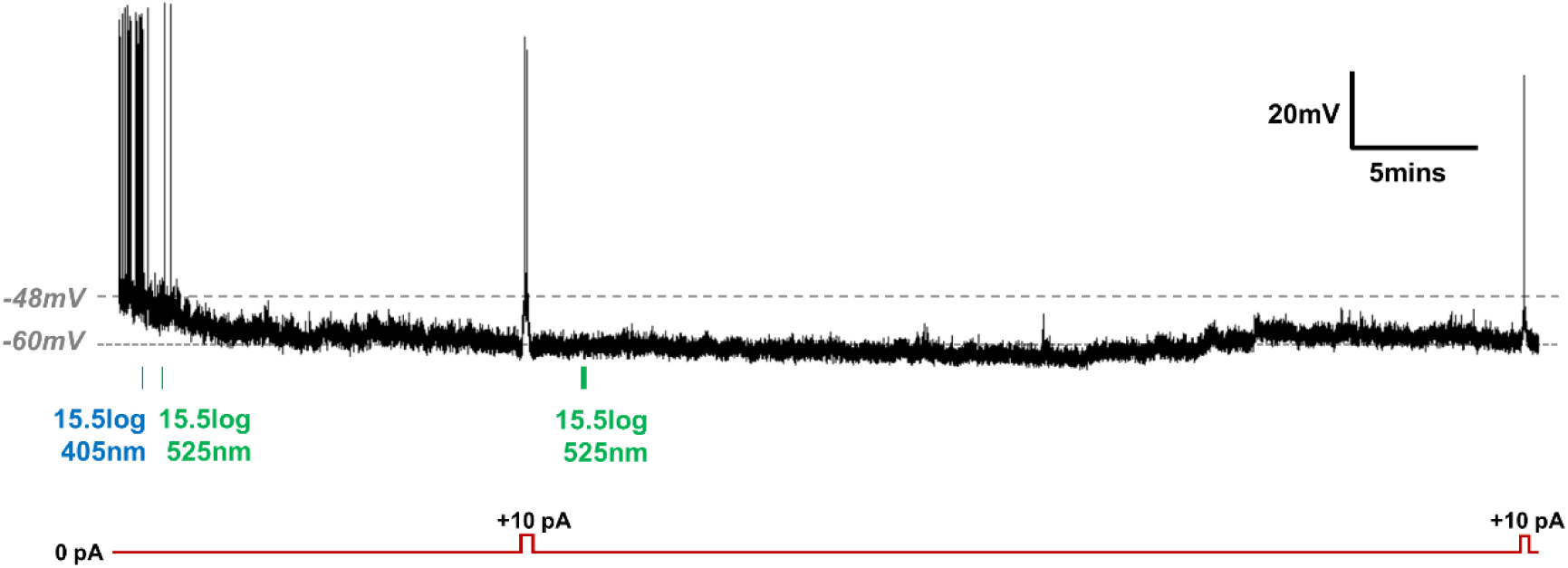
Some hyperpolarised SCN neurons do not respond to 525nm light. Example trace from a cell with prolonged inhibition. After exposure to bright 405nm light (15.5log photons/cm^2^/s, blue bar), this cell remained hyperpolarised for at least 55 minutes. This long-lasting hyperpolarisation was not due to issues with cell health as action potentials could still be elicited when we manually applied a depolarising current. In this cell, there was no obvious impact of 525nm light (15.5log photons/cm^2^/s, green bars). Schematic of depolarising pulse shown in red. We are not able to determine whether the failure of 525nm light to restore firing reflects some aspect of the Lamplight response in this cell or an unrelated change in spontaneous activity, as SCN neurons are known to switch between active and inactive states.

## Supplementary Table

**Table S1.** Ratiometric light stimuli used in Hek293T and retinal MEA experiments.

| Light Stimuli | Lamplight Inactive state<br>($\lambda_{\max} = 370\text{nm}$)<br>Effective Photons/cm<sup>2</sup>/s<br>(log photons) | Lamplight Active state<br>($\lambda_{\max} = 515\text{nm}$)<br>Effective Photons/cm<sup>2</sup>/s<br>(log photons) | Total Lamplight<br>Effective Photons/cm<sup>2</sup>/s<br>(log photons) |
| --- | --- | --- | --- |
| **Hek293T BRET Assay** |  |  |  |
| **405nm only** |  |  |  |
| *Fig 1D, E* | $1.93 \times 10^{15}$ (15.29log) | $1.24 \times 10^{15}$ (15.09log) | $3.15 \times 10^{15}$ (15.50log) |
| *Fig 1F* | $4.83 \times 10^{14}$ (14.68log) | $3.30 \times 10^{14}$ (14.52log) | $8.12 \times 10^{14}$ (14.91log) |
| **525nm only** |  |  |  |
| *Fig 1D, E* | $1.50 \times 10^{12}$ (12.18log) | $3.16 \times 10^{15}$ (15.50log) | $3.20 \times 10^{15}$ (15.50log) |
| *Fig 1E* | $6.43 \times 10^{11}$ (11.81log) | $1.58 \times 10^{15}$ (15.20log) | $1.58 \times 10^{15}$ (15.20log) |
| *Fig 1F* | $1.13 \times 10^{12}$ (12.05log) | $2.37 \times 10^{15}$ (15.37log) | $2.37 \times 10^{15}$ (15.37log) |
| **Ratiometric stimuli (405nm : 525nm)** |  |  |  |
| *Fig 1H, 1I, 1J* |  |  |  |
| 1 : 0 | $3.19 \times 10^{15}$ (15.50log) | $2.11 \times 10^{15}$ (15.32log) | $5.30 \times 10^{15}$ (15.72log) |
| 0.8 : 0.2 | $2.58 \times 10^{15}$ (15.41log) | $2.76 \times 10^{15}$ (15.44log) | $5.34 \times 10^{15}$ (15.73log) |
| 0.6 : 0.4 | $1.93 \times 10^{15}$ (15.28log) | $3.32 \times 10^{15}$ (15.52log) | $5.24 \times 10^{15}$ (15.72log) |
| 0.4 : 0.6 | $1.28 \times 10^{15}$ (15.11log) | $3.97 \times 10^{15}$ (15.60log) | $5.25 \times 10^{15}$ (15.72log) |
| 0.2 : 0.8 | $6.71 \times 10^{14}$ (14.83log) | $4.59 \times 10^{15}$ (15.66log) | $5.26 \times 10^{15}$ (15.72log) |
| 0 : 1 | $7.21 \times 10^{12}$ (12.86 log) | $5.19 \times 10^{15}$ (15.72log) | $5.20 \times 10^{15}$ (15.72log) |
| **Retinal MEA recordings** |  |  |  |
| **405nm only** |  |  |  |
| *Fig 3B-3F* | $6.13 \times 10^{15}$ (15.78log) | $4.08 \times 10^{15}$ (15.61log) | $1.02 \times 10^{16}$ (16.01log) |
| **525nm only** |  |  |  |
| *Fig 3B-3F* | $3.96 \times 10^{12}$ (12.59log) | $1.02 \times 10^{16}$ (16.01log) | $1.02 \times 10^{16}$ (16.01log) |
| **Ratiometric stimuli (405nm: 525nm)** |  |  |  |
| *Fig 3E, 3F* | $3.06 \times 10^{15}$ (15.49log) | $7.10 \times 10^{15}$ (15.85log) | $1.02 \times 10^{16}$ (16.01log) |

## References

1. Ferenczi, E. A., Tan, X. & Huang, C. L.-H. Principles of Optogenetic Methods and Their Application to Cardiac Experimental Systems. Front. Physiol. 10, (2019).

2. Wiegert, J. S., Mahn, M., Prigge, M., Printz, Y. & Yizhar, O. Silencing Neurons: Tools, Applications, and Experimental Constraints. Neuron 95, 504–529 (2017).

3. Han, X. & Boyden, E. S. Multiple-Color Optical Activation, Silencing, and Desynchronization of Neural Activity, with Single-Spike Temporal Resolution. PLOS ONE 2, e299 (2007).

4. Zhang, F. et al. Multimodal fast optical interrogation of neural circuitry. Nature 446, 633–639 (2007).

5. Chow, B. Y. et al. High-performance genetically targetable optical neural silencing by light-driven proton pumps. Nature 463, 98–102 (2010).

6. Chuong, A. S. et al. Noninvasive optical inhibition with a red-shifted microbial rhodopsin. Nat. Neurosci. 17, 1123–1129 (2014).

7. Govorunova, E. G., Sineshchekov, O. A., Janz, R., Liu, X. & Spudich, J. L. Natural light-gated anion channels: A family of microbial rhodopsins for advanced optogenetics. Science 349, 647–650 (2015).

8. Berndt, A., Lee, S. Y., Ramakrishnan, C. & Deisseroth, K. Structure-Guided Transformation of Channelrhodopsin into a Light-Activated Chloride Channel. Science 344, 420–424 (2014).

9. Alberio, L. et al. A light-gated potassium channel for sustained neuronal inhibition. Nat. Methods 15, 969–976 (2018).

10. Cosentino, C. et al. Optogenetics. Engineering of a light-gated potassium channel. Science 348, 707–710 (2015).

11. Ferenczi, E. & Deisseroth, K. When the electricity (and the lights) go out: transient changes in excitability. Nat. Neurosci. 15, 1058–1060 (2012).

12. Mattis, J. et al. Principles for applying optogenetic tools derived from direct comparative analysis of microbial opsins. Nat. Methods 9, 159–172 (2011).

13. Mahn, M., Prigge, M., Ron, S., Levy, R. & Yizhar, O. Biophysical constraints of optogenetic inhibition at presynaptic terminals. Nat. Neurosci. 19, 554–556 (2016).

14. Malyshev, A. Y. et al. Chloride conducting light activated channel GtACR2 can produce both cessation of firing and generation of action potentials in cortical neurons in response to light. Neurosci. Lett. 640, 76–80 (2017).

15. Armbruster, B. N., Li, X., Pausch, M. H., Herlitze, S. & Roth, B. L. Evolving the lock to fit the key to create a family of G protein-coupled receptors potently activated by an inert ligand. Proc. Natl. Acad. Sci. U. S. A. 104, 5163–5168 (2007).

16. Wettschureck, N. & Offermanns, S. Mammalian G proteins and their cell type specific functions. Physiol. Rev. 85, 1159–1204 (2005).

17. Urban, D. J. & Roth, B. L. DREADDs (Designer Receptors Exclusively Activated by Designer Drugs): Chemogenetic Tools with Therapeutic Utility. Annu. Rev. Pharmacol. Toxicol. 55, 399–417 (2015).

18. Whissell, P. D., Tohyama, S. & Martin, L. J. The Use of DREADDs to Deconstruct Behavior. Front. Genet. 7, (2016).

19. Lüscher, C., Jan, L. Y., Stoffel, M., Malenka, R. C. & Nicoll, R. A. G Protein-Coupled Inwardly Rectifying K+ Channels (GIRKs) Mediate Postsynaptic but Not Presynaptic Transmitter Actions in Hippocampal Neurons. Neuron 19, 687–695 (1997).

20. Stachniak, T. J., Ghosh, A. & Sternson, S. M. Chemogenetic Synaptic Silencing of Neural Circuits Localizes a Hypothalamus→Midbrain Pathway for Feeding Behavior. Neuron 82, 797–808 (2014).

21. Berry, M. H. et al. Restoration of high-sensitivity and adapting vision with a cone opsin. Nat. Commun. 10, 1–12 (2019).

22. Cehajic-Kapetanovic, J. et al. Restoration of Vision with Ectopic Expression of Human Rod Opsin. Curr. Biol. CB 25, 2111–2122 (2015).

23. Gaub, B. M., Berry, M. H., Holt, A. E., Isacoff, E. Y. & Flannery, J. G. Optogenetic Vision Restoration Using Rhodopsin for Enhanced Sensitivity. Mol. Ther. J. Am. Soc. Gene Ther. 23, 1562–1571 (2015).

24. Masseck, O. A. et al. Vertebrate Cone Opsins Enable Sustained and Highly Sensitive Rapid Control of Gi/o Signaling in Anxiety Circuitry. Neuron 81, 1263–1273 (2014).

25. Airan, R. D., Thompson, K. R., Fenno, L. E., Bernstein, H. & Deisseroth, K. Temporally precise in vivo control of intracellular signalling. Nature 458, 1025–1029 (2009).

26. Kawano-Yamashita, E. et al. Activation of Transducin by Bistable Pigment Parapinopsin in the Pineal Organ of Lower Vertebrates. PloS One 10, e0141280 (2015).

27. Eickelbeck, D. et al. Lamprey Parapinopsin (“UVLamP”): a Bistable UV-Sensitive Optogenetic Switch for Ultrafast Control of GPCR Pathways. ChemBioChem 21, 612–617 (2020).

28. Koyanagi, M. et al. Bistable UV pigment in the lamprey pineal. Proc. Natl. Acad. Sci. U. S. A. 101, 6687–6691 (2004).

29. Wada, S. et al. Color opponency with a single kind of bistable opsin in the zebrafish pineal organ. Proc. Natl. Acad. Sci. U. S. A. 115, 11310–11315 (2018).

30. Jiang, M. & Bajpayee, N. S. Molecular Mechanisms of Go Signaling. Neurosignals 17, 23–41 (2009).

31. Masuho, I. et al. Distinct profiles of functional discrimination among G proteins determine the actions of G protein-coupled receptors. Sci. Signal. 8, ra123 (2015).

32. Green, D. J. & Gillette, R. Circadian rhythm of firing rate recorded from single cells in the rat suprachiasmatic brain slice. Brain Res. 245, 198–200 (1982).

33. Hermanstyne, T. O., Simms, C. L., Carrasquillo, Y., Herzog, E. D. & Nerbonne, J. M. Distinct Firing Properties of Vasoactive Intestinal Peptide-Expressing Neurons in the Suprachiasmatic Nucleus. J. Biol. Rhythms 31, 57–67 (2016).

34. Inouye, S. T. & Kawamura, H. Persistence of circadian rhythmicity in a mammalian hypothalamic ‘island’ containing the suprachiasmatic nucleus. Proc. Natl. Acad. Sci. 76, 5962–5966 (1979).

35. Meijer, J. H., Watanabe, K., Détàri, L. & Schaap, J. Circadian rhythm in light response in suprachiasmatic nucleus neurons of freely moving rats. Brain Res. 741, 352–355 (1996).

36. Paul, S. et al. Output from VIP cells of the mammalian central clock regulates daily physiological rhythms. Nat. Commun. 11, 1–14 (2020).

37. Yamazaki, S., Kerbeshian, M. C., Hocker, C. G., Block, G. D. & Menaker, M. Rhythmic Properties of the Hamster Suprachiasmatic NucleusIn Vivo. J. Neurosci. 18, 10709–10723 (1998).

38. Dhingra, A. et al. The Light Response of ON Bipolar Neurons Requires Gαo. J. Neurosci. 20, 9053–9058 (2000).

39. Ivanova, E., Hwang, G.-S. & Pan, Z.-H. Characterization of transgenic mouse lines expressing Cre-recombinase in the retina. Neuroscience 165, 233–243 (2010).

40. Chang, B. et al. Retinal degeneration mutants in the mouse. Vision Res. 42, 517–525 (2002).

41. Pittler, S. J. & Baehr, W. Identification of a nonsense mutation in the rod photoreceptor cGMP phosphodiesterase beta-subunit gene of the rd mouse. Proc. Natl. Acad. Sci. U. S. A. 88, 8322–8326 (1991).

42. Eleftheriou, C. G. et al. Melanopsin Driven Light Responses Across a Large Fraction of Retinal Ganglion Cells in a Dystrophic Retina. Front. Neurosci. 14, (2020).

43. Tu, D. C. et al. Physiologic Diversity and Development of Intrinsically Photosensitive Retinal Ganglion Cells. Neuron 48, 987–999 (2005).

44. Weng, S., Estevez, M. E. & Berson, D. M. Mouse ganglion-cell photoreceptors are driven by the most sensitive rod pathway and by both types of cones. PloS One 8, e66480 (2013).

45. Syrovatkina, V., Alegre, K. O., Dey, R. & Huang, X.-Y. Regulation, Signaling, and Physiological Functions of G-Proteins. J. Mol. Biol. 428, 3850–3868 (2016).

46. Lüscher, C. & Slesinger, P. A. Emerging roles for G protein-gated inwardly rectifying potassium (GIRK) channels in health and disease. Nat. Rev. Neurosci. 11, 301–315 (2010).

47. Hauser, A. S., Attwood, M. M., Rask-Andersen, M., Schiöth, H. B. & Gloriam, D. E. Trends in GPCR drug discovery: new agents, targets and indications. Nat. Rev. Drug Discov. 16, 829–842 (2017).

48. Roth, B. L. DREADDs for Neuroscientists. Neuron 89, 683–694 (2016).

49. Tichy, A.-M., Gerrard, E. J., Sexton, P. M. & Janovjak, H. Light-activated chimeric GPCRs: limitations and opportunities. Curr. Opin. Struct. Biol. 57, 196–203 (2019).

50. Xu, Y. et al. The TRPM1 channel in ON-bipolar cells is gated by both the α and the βγ subunits of the G-protein Go. Sci. Rep. 6, 20940 (2016).

51. Marino, S. et al. PTEN is essential for cell migration but not for fate determination and tumourigenesis in the cerebellum. Dev. Camb. Engl. 129, 3513–3522 (2002).

52. Peachey, N. S. et al. GPR179 is required for depolarizing bipolar cell function and is mutated in autosomal-recessive complete congenital stationary night blindness. Am. J. Hum. Genet. 90, 331–339 (2012).

53. Taniguchi, H. et al. A resource of Cre driver lines for genetic targeting of GABAergic neurons in cerebral cortex. Neuron 71, 995–1013 (2011).

54. Petrs-Silva, H. et al. Novel properties of tyrosine-mutant AAV2 vectors in the mouse retina. Mol. Ther. J. Am. Soc. Gene Ther. 19, 293–301 (2011).

55. Belle, M. D. C. & Piggins, H. D. Circadian regulation of mouse suprachiasmatic nuclei neuronal states shapes responses to orexin. Eur. J. Neurosci. 45, 723–732 (2017).

56. Hanna, L., Walmsley, L., Pienaar, A., Howarth, M. & Brown, T. M. Geniculohypothalamic GABAergic projections gate suprachiasmatic nucleus responses to retinal input. J. Physiol. 595, 3621–3649 (2017).

57. Egert, U. & Meyer, T. Heart on a Chip — Extracellular Multielectrode Recordings from Cardiac Myocytes in Vitro. in Practical Methods in Cardiovascular Research (eds. Dhein, S., Mohr, F. W. & Delmar, M.) 432–453 (Springer Berlin Heidelberg, 2005). doi: 10.1007/3-540-26574-0_22.

58. Lelong, I. H., Petegnief, V. & Rebel, G. Neuronal cells mature faster on polyethyleneimine coated plates than on polylysine coated plates. J. Neurosci. Res. 32, 562–568 (1992).

59. Govardovskii, V. I., Fyhrquist, N., Reuter, T., Kuzmin, D. G. & Donner, K. In search of the visual pigment template. Vis. Neurosci. 17, 509–528 (2000).

